# DRUID: A pipeline for reproducible transcriptome-wide measurements of mRNA stability

**DOI:** 10.1101/149195

**Authors:** Andrew Lugowski, Beth Nicholson, Olivia S. Rissland

**Affiliations:** Molecular Medicine Program, The Hospital for Sick Children Research Institute, Toronto, ON M5G 0A4, Canada; Department of Molecular Genetics, University of Toronto, Toronto, ON M5S 1A8 Canada

## Abstract

Control of messenger RNA (mRNA) stability is an important aspect of gene regulation. The gold standard for measuring mRNA stability transcriptome-wide uses metabolic labeling, biochemical isolation of labeled RNA populations, and high-throughput sequencing. However, difficult normalization procedures and complex computational algorithms have inhibited widespread adoption of this approach. Here, we present DRUID (for Determination of Rates Using Intron Dynamics), a computational pipeline that is robust, easy to use, and freely available. Our pipeline uses endogenous introns to normalize time course data and yields more reproducible half-lives than other methods, even with datasets that were otherwise unusable. DRUID can handle datasets from a variety of organisms, spanning yeast to humans, and we even applied it retroactively on published datasets. We anticipate that DRUID will allow broad application of metabolic labeling for studies of mRNA stability.

## INTRODUCTION

A critical component in controlling gene expression, RNA decay is essential for nearly all biological processes, from early development to inflammatory responses (Giraldez et al. 2006; Tadros et al. 2007; Brooks and Blackshear 2013). However, transcriptome-wide measurements of mRNA half-lives have long been challenging and represent a major barrier for broad investigations of how mRNA stability is regulated. One strategy has been to shut off transcription, thereby repressing synthesis of all transcripts. In *Saccharomyces cerevisiae*, these experiments typically involve using a temperature-sensitive mutant of RNA polymerase II (Herrick et al. 1990; Grigull et al. 2004; Presnyak et al. 2015), and work in mammalian cell lines has predominantly relied upon drugs that target RNA polymerases, such as actinomycin D and α-amanitin (Ross 1995; Bensaude 2011). Each of these treatments places substantial stress on the cell and can alter the stability and localization of numerous transcripts, as well as broader phenotypes like cell growth (Bensaude 2011; Sun et al. 2013; Geisberg et al. 2014).

Metabolic labeling has emerged as a powerful alternative strategy for determining RNA stabilities under more physiological conditions (Ross 1995; Rabani et al. 2011; Tani et al. 2012; Neymotin et al. 2014; Duffy et al. 2015). This approach uses nucleobase or nucleoside analogs, such as 4-thiouridine (4SU), 5-bromouridine (BrU), or 5-ethynyl uridine (EU), all of which allow subsequent isolation of labeled RNA populations (Rabani et al. 2011; Tani et al. 2012; Imamachi et al. 2014; Neymotin et al. 2014; Duffy et al. 2015). The selected RNA is then quantified, often by high-throughput sequencing.

Several experimental variations of metabolic-labeling methods have been described (Munchel et al. 2011; Tani et al. 2012; Imamachi et al. 2014; Neymotin et al. 2014). One common method uses an approach-to-equilibrium strategy (Figure 1A). Here, cells are harvested after increasing times of incubation with the nucleoside analog. Over the time course, mRNA abundance of the labeled population approaches steady-state levels. Because the rate of this approach is determined by transcript stability, these measurements can be used to infer half-lives (Greenberg 1972; Ross 1995; Neymotin et al. 2014). However, the resultant data are complicated and not easily analyzed, unlike shut-off experiments whose results can be fit to simple exponential decay models. Thus, despite its clear benefits, the approach-to-equilibrium method remains inaccessible to the general researcher, and polymerase inhibitors remain much more broadly used (Burow et al. 2015; Kumagai et al. 2016; Mauer et al. 2016; Ayupe and Reis 2017).

**Figure 1.**
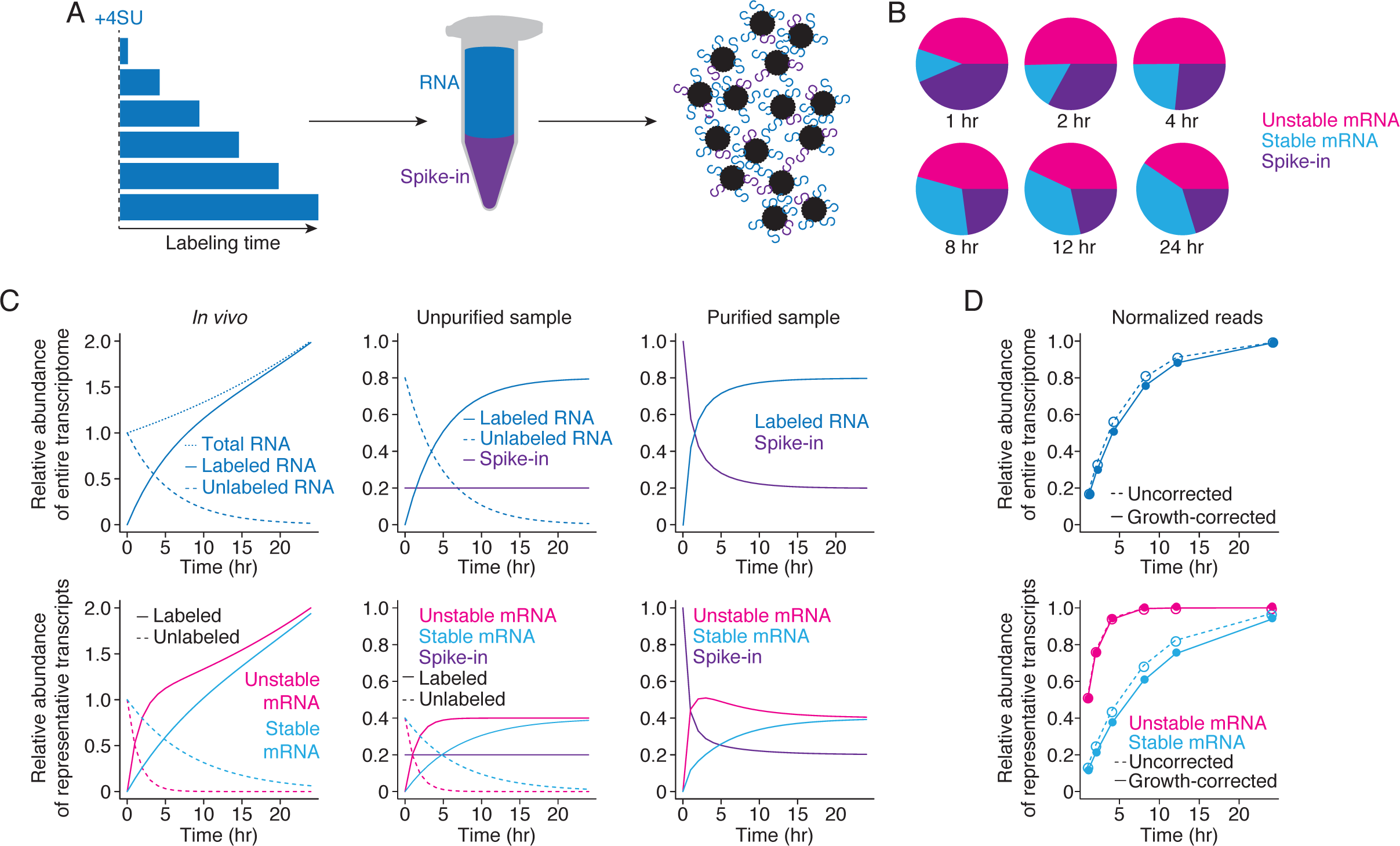
Conceptual underpinnings of the approach-to-equilibrium method. (A) Representation of the approach-to-equilibrium experimental method. Cells are incubated with 4SU for increasing amounts of time. Isolated RNA is then biotinylated *in vitro* and purified using streptavidin. The eluate is then prepared for high-throughput RNA sequencing. (B) Representation of typical RNA-seq results. Reads for each gene represent a fraction of the total library, which changes through time, depending on individual synthesis and decay rates as well as overall transcription levels. Representative results are shown for an unstable transcript (pink), a stable transcript (blue), and a spike-in (purple). (C) Simulation of abundances during an approach-to-equilibrium experiment for the transcriptome (top) and individual transcripts (bottom). Left: As cells are incubated longer with 4SU, the labeled RNA population increases at a rate sufficient to replace the unlabeled population and for cell growth. The unlabeled population for unstable transcripts (pink) necessarily decreases faster than for stable transcripts (blue). Center: Once the RNA is extracted, spike-ins are added at a constant ratio to the total amount of RNA, and information about cell growth is lost. Right: After purification, the ratio of spike-in to labeled sample behaves stereotypically with a rapid decrease over the initial time points and convergence to an asymptote, which represents the ratio of the RNAs in the unpurified sample. Note that at early time points, unstable transcripts represent a larger fraction of the library than their eventual steady-state ratios. (D) Simulation of normalization strategies. The pie charts (from (B)) are normalized to spike-in levels to calculate the relative abundance of the entire transcriptome or specific transcripts, as appropriate. These measurements can then be growth-corrected, and, using a bounded growth equation, half-lives can be calculated. Note that correcting for growth has a larger effect on stable transcripts than on unstable ones.

Here, we describe DRUID, a computational method for determining mRNA half-lives on a transcriptome-wide scale using metabolic labeling. By normalizing to intron-mapping reads, our method allowed us to determine mRNA stabilities with higher reproducibility and ease than other methods. DRUID can also rescue poorly behaving, and previously unusable, datasets, and can even be applied to datasets from species with few introns, such as *S. cerevisiae*. Finally, we have developed a computational package that is publically available to enable broader use by the community.

## DESIGN

### The conceptual underpinnings of metabolic labeling and approach-to-equilibrium kinetics

The approach-to-equilibrium strategy relies upon incorporation of a nucleotide analog into nascent transcripts so that, with increasing incubation periods, the fraction of transcripts labeled increases until steady-state has been reached. The RNA-seq readout of metabolic-labeling experiments can be envisioned as a series of pie-charts through time with each slice representing the relative proportion of reads mapping to each transcript (Figure 1B). Unlike most RNA-seq experiments, where only a handful of transcripts will change in abundance, during metabolic labeling experiments, the relative and absolute abundance of every transcript will change over the experiment (Figure 1C). For an individual transcript, its relative abundance is determined by synthesis and decay rates, but the absolute abundance of the total labeled population increases with longer exposure to the nucleotide analog at a rate sufficient to replace degraded, unlabeled RNA as well as to allow for cell growth (Figure 1C, left panels). Determining mRNA half-lives requires taking both classes of behaviors into account.

The current solution uses exogenously added spike-ins to convert relative RNA-seq measurements into absolute measurements of RNA abundance (Figure 1C, middle panels). Typically, spike-ins are added to the harvested RNA from each time point prior to biochemical purification, and information about growth rate is necessarily lost at this step. Because the proportion of spike-ins to the total RNA sample remains constant, once the labeled population has been purified *in vitro*, the proportion of spike-ins decreases over time while that of the labeled sample increases. These ratios approach that in the original, unpurified sample (Figure 1C, right panels). Individual transcripts will reach equilibrium with differing kinetics, determined entirely by their stability, with the most unstable transcripts reaching equilibrium the fastest. To calculate half-lives, the RNA-seq libraries are normalized to the spike-ins and further corrected for cell growth. A bounded growth equation is used to infer the decay rate of the unlabeled RNA population (Figure 1D).

Spike-ins are thus absolutely essential for determining half-lives using metabolic labeling. Although there are commercially available RNA-seq spike-ins (such as the ERCC set), these lack nucleotide analogs, such as 4SU, and so currently each laboratory *in vitro* transcribes their own standards, as needed (Imamachi et al. 2014; Neymotin et al. 2014). However, there is no broadly accepted standard for these spike-ins, in terms of the number of transcripts, their nucleotide make-up, or length, and variations in spike-in make-up can have large effects on the eventual half-life determination. Indeed, previous attempts using three *in vitro* transcribed spike-ins were associated with enough technical noise that biological replicates were combined in order to determine half-lives (Neymotin et al. 2014).

### DRUID: motivation and advantages

We initially identified two major barriers for the acceptance of the approach-to-equilibrium method: the cost, difficulty, and technical variation associated with current spike-ins; and the computational complexities for analyzing subsequent datasets. Our goal was to develop a method easy enough for widespread use of metabolic labeling, and so we set about reducing these barriers.

Because of the variability inherent in using a handful of *in vitro* transcribed spike-ins, we initially opted to use RNA from two organisms that differed from the species being interrogated. For example, in experiments with HEK293 cells, we used 4SU-labeled RNA from *Drosophila* S2 cells (for normalization) and unlabeled *S. cerevisiae* RNA (to determine the enrichment obtained during the purification). These spike-ins were easy and cheap to generate, produced reproducible half-life measurements (see below), and were easily applied to mouse and human cell lines.

In the process of optimizing this method, we produced datasets that did not contain sufficient amounts of spike-ins for robust normalization. We asked whether normalization to some internal RNA species could salvage these datasets. Because introns are highly unstable, they reach equilibrium very quickly and can thus serve as a normalization factor. We thus developed DRUID, a robust method that normalizes to intron-mapping reads and calculates half-lives. In contrast to other methods and our original design, DRUID can be applied retroactively to datasets not originally designed for this scheme. DRUID half-life measurements also show higher reproducibility than other methods, most likely because introns provide additional controls for a range of technical variables, such as 4SU-labeling efficiency and cell growth.

## RESULTS

### Normalizing to exogenous whole organism spike-ins allows for the calculation of RNA half-lives

To determine mRNA half-lives in HEK293 cells, we initially used two sets of spike-ins: 4SU-labeled RNA from *Drosophila* S2 cells and unlabeled RNA from *S. cerevisiae*. The *Drosophila* and *S. cerevisiae* genomes differ sufficiently enough from the human one that even short reads can be unambiguously assigned (Figure S1A). We routinely observed 100-fold enrichment of 4SU-labeled *Drosophila* RNA compared to unlabeled yeast RNA, indicating that less than 1% of the signal in the selected population was due to nonspecific background (Figure 2A). Subsequent analysis indicated that the S2 RNA was less labeled than the HEK293 RNA (Figure S1B, C), and so these enrichment values likely represent a lower bound.

**Figure 2.**
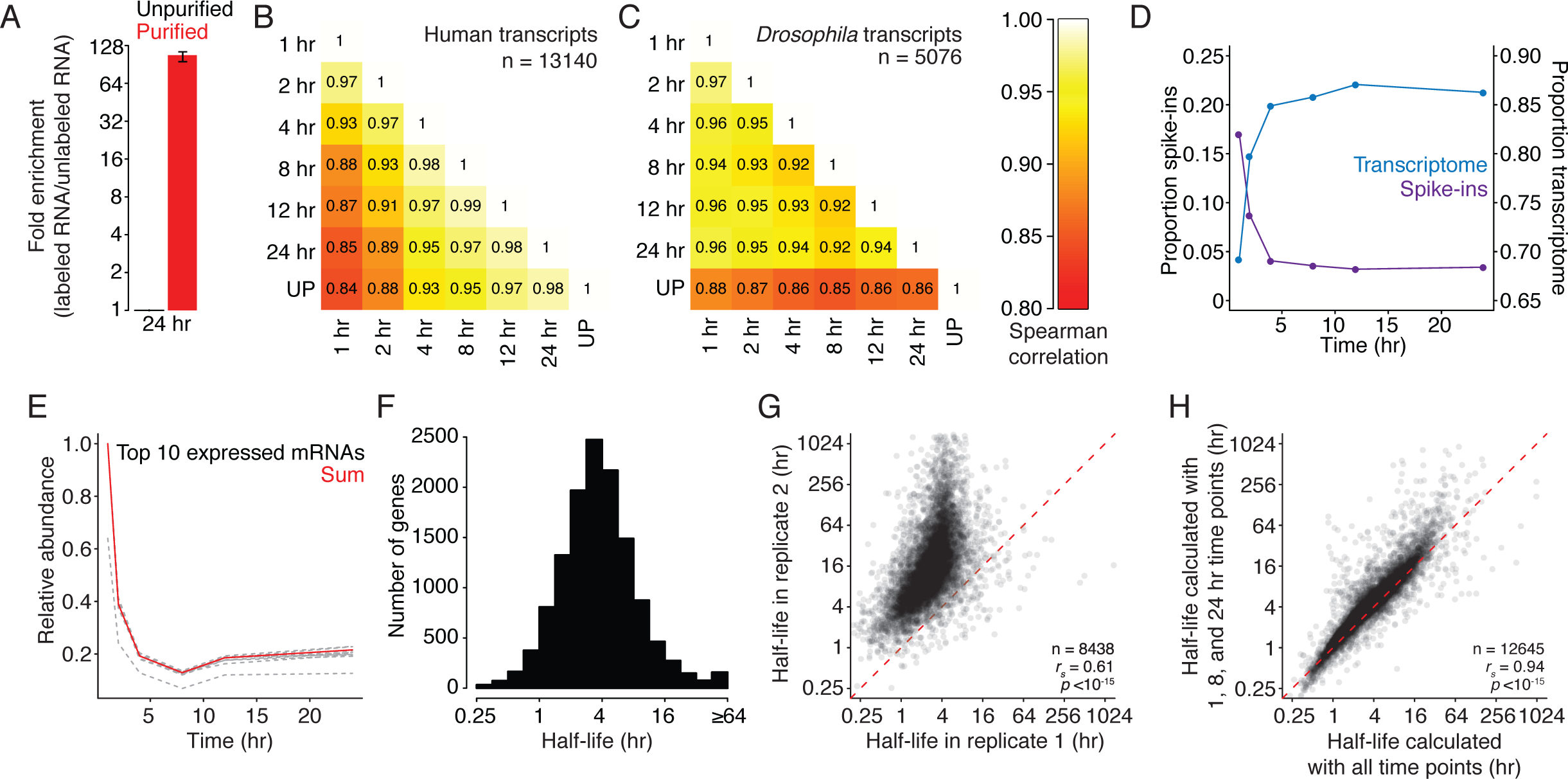
Exogenous, whole-organism spike-ins can be used to calculate mRNA half-lives. (A) Enrichment of 4SU-labeled transcripts. Plotted is the mean enrichment of S2 RNA compared to yeast RNA for purified 24-hr samples (red) relative to unpurified (black) samples for two biological replicates. The black line represents range. (B) Comparisons of human mRNA abundance between samples. A heatmap is plotted comparing the abundance of human transcripts in each purified sample, as well as the unpurified (UP) sample. Values in each box are Spearman correlations. (C) Comparisons of fly mRNA abundances between samples. As in (B), except for transcripts encoded in the *Drosophila* genome. (D) Ratios of fly to human reads through the time course. The fraction of reads mapping to the human (blue) or *Drosophila* (purple) genome is plotted for each time point. (E) The behavior of individual *Drosophila* genes. Plotted is relative abundance for each of the top 10 most highly expressed genes (dashed lines) and the sum of reads mapping to these genes (in red). (F) Distribution of mRNA half-lives in human cells. The histogram shows the distribution of mRNA half-lives in human cells for 12890 genes. (G) Comparison of mRNA half-lives between biological replicates. A scatterplot comparing human mRNA half-lives for two biological replicates. The red dashed line represents the x = y line. (H) mRNA half-lives calculated using only three time points. A scatterplot comparing mRNA half-lives calculated by using all time points or only three (1, 8, and 24 hr). The red dashed line represents the x = y line.

As incubation with 4SU increased, samples showed reduced similarity in the abundance of human mRNAs (Figure 2B). Consistent with most transcripts having reached equilibrium by the last time point, the unpurified sample was least similar to the 1-hr sample and most similar to the 24 hr one (*r_s_* = 0.84 and *r_s_* = 0.98, respectively). In contrast, reads mapping to the *Drosophila* genomes were unaffected by different harvesting times (Figure 2C; *r_s_* = 0.92 to 0.97).

Longer 4SU-incubation times also resulted in a higher fraction of reads mapping to the human genome and a lower fraction to the *Drosophila* genome (Figure 2D). Although the behavior of individual transcripts from the S2-cell spike-in varied, the sum of all reads mapping to the *Drosophila* genome was resistant to outliers (Figure 2E). We normalized the human-mapping reads by the sum of *Drosophila*-mapping reads, fit a bounded-growth equation to the data, and corrected for the average doubling time in HEK293 cells. Unlike other approaches (Dölken et al. 2008), we did not rely upon the unselected sample for calculating half-lives, and thus our calculations were unaffected by differential selection biases (such as that reported for longer transcripts [Duffy et al. 2015]).

We calculated half-lives for 12,890 genes in HEK293 cells (Figure 2F). There was wide variation in mRNA stability in these cells, with the most unstable transcript (ID1) having a half-life less than 15 minutes, and others having half-lives longer than the cell cycle. For these long-lived transcripts, cell growth and dilution make a major contribution to their dynamics. The half-lives were similar to those we generated with actinomycin D and α-amanitin (Figure S1E, F; *r_s_* = 0.65 and 0.62, respectively) and with those previously reported (*r_s_* = 0.62 [Tani et al. 2012]). These half-lives were generally reproducible (Figure 2G; *r_s_* = 0.61), although we noted that the longest-lived genes varied the most in magnitude (but not rank order) between replicates.

In these experiments, we had included six time points in addition to steady-state measurements. However, we hypothesized that not all of these time points would be necessary to determine stabilities, and using fewer samples could reduce the associated library preparation and sequencing costs. We repeated our analysis, but this time omitting individual time points. With the exception of the 24 hr time point, the calculated half-lives were robust to the omission of a single time point (Figure S1F; *r_s_* = 0.95 to 1 vs. *r_s_* = 0.89). The 24-hr time point is likely critical for calculating half-lives, especially for more stable transcripts, because it represents near-equilibrium measurements. Surprisingly, using only three time points (1, 8, and 24 hr) gave similar measurements as with all the time points (Figure 1H; *r_s_* = 0.94). Although calculations become more robust with more time points, these three time points represent the minimum requirement for calculating half-lives in human cells using the approach-to-equilibrium method.

### Internal short-lived RNA species can be used to determine mRNA stabilities

In the process of obtaining these half-life measurements, we generated flawed datasets that resulted from insufficient spike-in reads or inconsistent labeling bias, each of which gave a characteristic signature (Figure S2A, B). Such datasets would typically be excluded from downstream analysis, but we wondered if they could be rescued with alternative normalization methods. We hypothesized that because unstable endogenous RNA species quickly reach equilibrium, they could perform a role similar to that of the labeled spike-in RNA.

We noted that introns were especially abundant at the earliest time points, making up 19% of the total reads (Figure 3A). Moreover, as with the *Drosophila* spike-ins, the overall proportion of reads mapping to introns exhibited a time-dependent decrease (Figure 3A), and the relative abundance of individual introns did not show a large time-dependent decrease in similarity (Figure S2C; *r_s_* = 0.90 to 0.95), indicating that equilibrium levels were generally reached before the first time point. However, we noted that not all introns behave similarly (Figure 3B). While 1045 introns showed a time-dependent decrease in levels, 1564 introns, although expressed at sufficient levels, did not show evidence of being highly unstable. Unexpectedly long-lived introns were likely included in the eventual mature transcript, either due to intron retention or alternative splicing (Braunschweig et al. 2014), or were misannotated.

**Figure 3.**
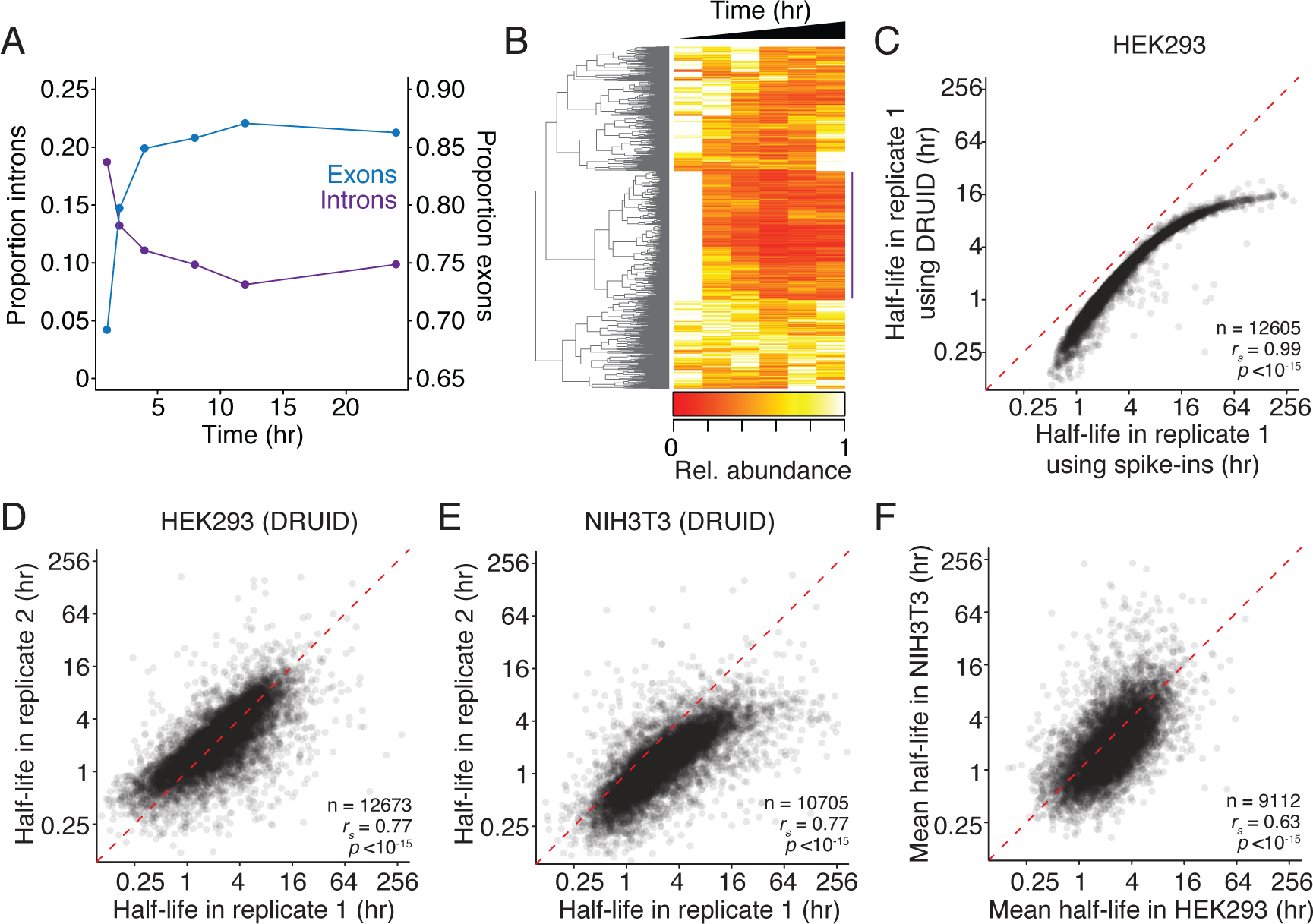
DRUID uses intron measurements to calculate mRNA half-lives. (A) Behavior of reads mapping to introns during the 4SU-labeling time course. The fraction of reads mapping to exons (blue) or introns (purple) is plotted for each time point in replicate 1. (B) Dynamics of intron abundance. Individual introns expressed in HEK293 cells were clustered by their behavior over the 4SU time course in replicate 1. Those introns showing the expected decrease (marked by the purple line) were used in downstream analyses. (C) Comparison of half-lives calculated using exogenous spike-ins or DRUID. A scatterplot comparing mRNA half-lives in HEK293 cells calculated with exogenous spike-ins or with introns (DRUID). The red dashed line represents the x = y line. (D) Reproducibility of human mRNA half-lives calculated using DRUID. A scatterplot comparing mRNA half-lives determined using DRUID for two biological replicates. The red dashed line represents the x = y line. (E) Reproducibility of mRNA half-lives in NIH3T3 cells calculated using DRUID, otherwise as in (D). (F) Comparison of HEK293 and NIH3T3 mRNA half-lives. A scatterplot comparing mean mRNA half-lives in HEK293 and NIH3T3 cells. The red dashed line represents the x = y line.

We used the sum of all reads mapping to the well-behaved introns and calculated half-lives for 12,673 genes in a pipeline we termed “DRUID”. Unlike with exogenous spike-ins, DRUID does not require correction for cellular growth. These half-lives correlated with those obtained by normalizing to exogenous spike-ins (Figure 3C; *r_s_* = 0.99), although they were slightly shorter. Surprisingly, normalization to introns yielded half-lives even more reproducible than those we obtained earlier (*r_s_* = 0.77 vs. *r_s_* = 0.61). Moreover, the skew that we had observed between replicates for long-lived transcripts was not apparent when we used DRUID (Figure 3D *cf*. Figure 2G), indicating that DRUID gives reproducible rank order and magnitudes for mRNA half-lives. Intron normalization likely captures *in vivo* experimental variation better than the exogenous spike-ins, and is thus better-equipped to normalize for these differences. When we generated half-lives using only three time points (1, 8, and 24 hr), DRUID gave similar results (Figure S2D; *r_s_* = 0.92) and was reproducible between replicates (*r_s_* = 0.73). Together, our analyses indicate that reproducible half-lives can be obtained by normalizing to introns and using only three time points.

### Orthologous mouse and human genes have similar mRNA half-lives

To further confirm the applicability of our method, we used DRUID to calculate mRNA half-lives in NIH3T3 cells using well-behaving introns. We obtained measurements for 10,705 genes with high similarity between replicates (Figure 3E; *r_s_* = 0.77). As with our HEK293 experiments, DRUID performed better than using exogenous spike-ins for normalization (Figure S2E; *r_s_* = 0.70). These values were similar to those previously calculated (Figure S2F; *r_s_* = 0.66, [Schwanhäusser et al. 2011]), although using our method we were able to determine half-lives for a much larger number of genes (5,029 vs. 10,705).

We next compared mRNA half-lives and equilibrium levels of orthologous human and mouse genes (Figure 3F, Figure S2G). Surprisingly, given that these two cell lines are from different organisms and derived from different cell types, we found a high correlation between RNA abundance and half-lives between mouse and human orthologs (*r_s_* = 0.57 and 0.63, respectively). Thus, although some transcripts, such as MBNL3, display striking differences in stability between HEK293 and NIH3T3 cells (26 hours vs. 36 minutes, respectively), many conserved transcripts are degraded at similar rates.

### DRUID can retroactively rescue datasets

Having established that DRUID can be used on high quality datasets, we finally asked whether this normalization method could rescue previously unusable datasets. We focused on two datasets: one with too few spike-in reads and the other with abnormal spike-in behavior (Figure S2A, B). For the first dataset, we were unable to calculate half-lives using the exogenous spike-ins. For the second, we obtained half-lives for only 3987 genes. Although these measurements did correlate with other replicates (Figure 4A, *r_s_* = 0.51), we observed a strong skew for long-lived transcripts.

**Figure 4.**
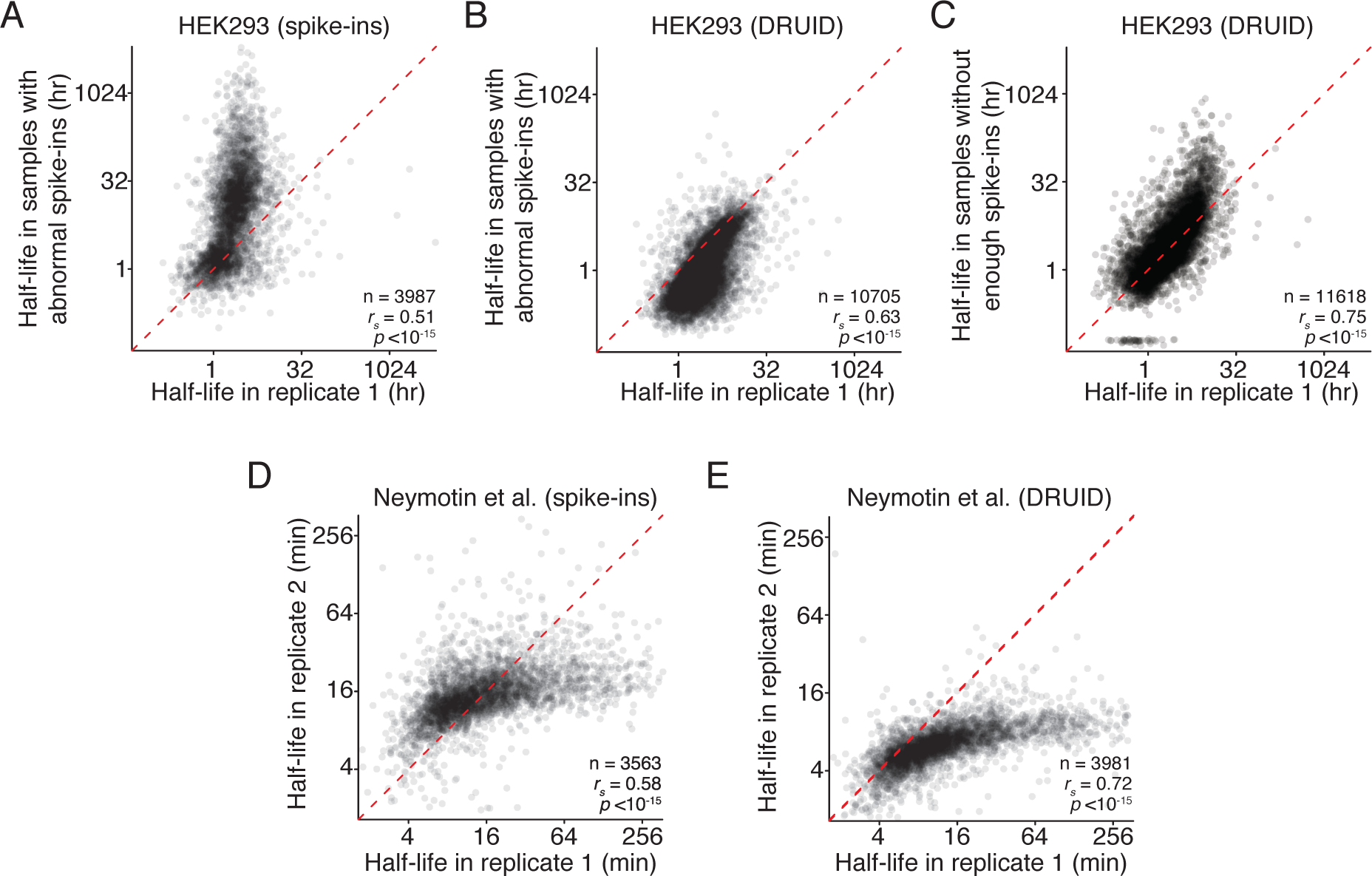
DRUID can rescue recalcitrant datasets. (A) Effect of poorly behaving datasets on mRNA half-lives calculated with exogenous spike-ins. A scatterplot comparing the half-lives in HEK293 cells calculated in one biological replicate and a recalcitrant dataset with poorly behaving spike-ins. The red dashed line represents the x = y line. (B) Comparison of mRNA half-lives calculated with DRUID. As in (A), except half-lives were calculated using DRUID. (C) Comparison of mRNA half-lives calculated with DRUID. As in (B), except for a second, recalcitrant dataset, whose spike-ins behaved so poorly that no half-lives could be calculated using exogenous spike-ins. (D) Comparison of yeast mRNA half-lives calculated using exogenous spike-ins, otherwise as in (A). (E) Comparisons of yeast mRNA half-lives calculated using DRUID, otherwise as in (A).

Strikingly, for both datasets, intron normalization was able to overcome both types of technical difficulties, and we generated half-lives for over 10,000 genes. These half-lives correlated well with our other datasets (Figure 4B, C; *r_s_* = 0.63–0.75). Importantly, we no longer observed the difference in half-life magnitude for stable transcripts that we saw with exogenous normalization (Figure 4A vs. Figure 4B). Thus, DRUID can be used for otherwise recalcitrant datasets.

One potential drawback of intron normalization is its applicability to organisms with few introns, such as *S. cerevisiae*. We thus applied DRUID to published datasets from budding yeast (Neymotin et al. 2014). When we calculated half-lives using the three exogenous spike-in transcripts, RNA half-lives were correlated (Figure 4D; *r_s_* = 0.58). However, DRUID generated half-lives that were better correlated (Figure 4E; *r_s_* = 0.72) and for a larger number of genes (3563 vs. 3981). We note that, irrespective of the computational scheme, the magnitude of half-lives differed between these two replicates, suggesting that there may be additional technical issues, such as labeling bias, that are independent of normalization schemes. These results indicate that normalizing to introns can be used on datasets not designed for DRUID. Furthermore, DRUID is a robust and widely applicable normalization method, appropriate even for organisms with few introns.

## DISCUSSION

Despite known and important issues in transcriptional shut-off approaches, RNA polymerase inhibitors remain in common use for determining transcript stability. To enable wider adoption of approach-to-equilibrium metabolic labeling strategies, we developed DRUID, a computational method that robustly calculates mRNA half-lives on a transcriptome-wide scale. Although we initially envisioned using exogenous spike-ins for a normalization approach, we were surprised that this framework was surpassed by intron normalization. DRUID was effective for all datasets we examined, irrespective of the organism examined, even those that were otherwise recalcitrant. Our computational pipeline is publically available to enable wider use of the approach-to-equilibrium strategy (see Methods).

Given the success of DRUID for calculating half-lives, is there any utility for including exogenous spike-ins? Although in principle they are not required, in practice, we still routinely include them in our experiments for two reasons. First, our exogenous spike-in strategy allows us to calculate the enrichment of labeled RNA in each dataset, thus confirming that the purification has worked as expected. Second, and more importantly, the exogenous spike-ins provide an independent normalization scheme and thus a useful quality control. Comparing between normalization schemes greatly increases confidence in the calculated half-lives and is particularly important when new cell types or systems are being used.

### Limitations

As with all approach-to-equilibrium strategies (Greenberg 1972; Ross 1995), there are two main requirements for DRUID. First, the labeling reagent, such as 4SU, must be readily taken up by the cell and incorporated into newly synthesized transcripts at concentrations that do not have negative physiological effects. Second, an underlying assumption of metabolic labeling and DRUID is that the system is at steady-state. Thus, in its current form, DRUID cannot be used to investigate scenarios where rates of synthesis and decay change throughout the experiment. Of course, biological processes, such as differentiation, are frequently defined by changes in both mRNA transcription and decay, and so an important next step is to generate experimental and computational methods that can monitor dynamic systems while remaining accessible to the broader community.

## ACKNOWLEDGEMENTS

We thank Dr. Andrew Spence, Dr. Julie Claycomb, and members of the Rissland and Claycomb labs for insightful questions and stimulating conversations. We especially thank Dr. Erik Sontheimer for his helpful feedback. We thank the Claycomb and Rubinstein labs for providing reagents. This work was funded by an NSERC Discovery Grant (to OSR), an Ontario Graduate Scholarship award (to AL), and an NSERC CGS-M award (to AL).

## METHODS

### CONTACT FOR REAGENT AND RESOURCE SHARING

Further information and requests for resources and reagents should be directed to and will be fulfilled by the corresponding author, Olivia Rissland.

### EXPERIMENTAL MODEL AND SUBJECT DETAILS

#### Cell lines

Human HEK293 epithelial cells (ATCC CRL1573) were cultured in Eagle’s Minimum Essential Medium (EMEM) supplemented with 10% Fetal Bovine Serum (FBS) and 1% penicillin-streptomycin solution. Murine NIH3T3 fibroblasts (ATCC CRL1658) were cultured in Dulbecco’s Modified Eagle’s Medium (DMEM) supplemented with 10% Donor Calf Serum (DCS) and 1% penicillin-streptomycin solution. Both mammalian cell lines were cultured at 37°C in a humidified incubator with 5% CO2. *Drosophila melanogaster* Schneider 2 (S2) cells (Thermo Fisher Scientific R69007) were cultured in ExpressFive SFM media (Thermo Fisher Scientific) supplemented with 10% heat-inactivated FBS and 20 mM L-Glutamine, at 28°C.

#### Organisms/Strains

*Saccharomyces cerevisiae* USY006 was grown in YPD liquid or plates at 30°C. RNA was isolated using the standard hot phenol method (Rissland and Norbury 2009). Synchronized populations of L1 *Caenorhabditis elegans* were grown on NGM plates for 60 hr until adult staged. Worms were washed off of plates with PBS buffer and resuspended in ultrapure water. RNA was extracted using TRI reagent (Molecular Research Center), according to manufacturer’s instructions.

### METHOD DETAILS

#### Metabolic labeling

For metabolic labeling experiments cells were treated with 100 μM 4SU and harvested after 1, 2, 4, 8, 12, and 24 hr. S2 cells treated with 100 μM 4SU for 24 hr were used for the generation of labeled spike-ins. When harvesting adherent cells, cells were dislodged with PBS and subjected to two PBS washes. S2 cells were pelleted and subjected to two PBS washes. RNA was extracted using TRI reagent (Molecular Research Center), according to manufacturer’s instructions.

#### Transcription shut-off experiments

HEK293 cells were treated with either 5 μg/mL actinomycin D or 50 μg/mL α-amanitin for 0, 1, 2, 4, 8, 12, and 24 hr and were harvested as described above.

#### *In vitro* biotinylation and biotin-streptavidin pull down

A 1 mg/mL solution of HPDP-biotin (Thermo Fisher Scientific) in dimethylformamide was incubated at 37°C for 30 min. 40-100 μg of RNA was combined with 20% w/w unlabeled yeast RNA and 20% w/w 4SU labeled fly RNA (human RNA for fly samples) and 120 μL of 2.5x citrate buffer (25 mM citrate pH 4.5, 2.5 mM EDTA) in a total volume of 240 μL. Sixty microliters of HPDP-biotin solution was added, and the RNA was incubated for 2 hr at 37°C, covered and shaking at 300 rpm. RNA was then phenol-chloroform extracted and ethanol precipitated with 2 μL of glycoblue (Life Technologies). The RNA pellet was resuspended in 200 μL of 1x wash buffer (10 mM Tris-Cl pH 7.4, 50 mM NaCl, 1 mM EDTA).

Fifty microliters of MACS microbeads (Miltenyi Biotec) were incubated with 48 μL of 1x wash buffer and 2 μL of yeast tRNA for 20 min at room temperature with rotation.

MACS microcolumns (Miltenyi Biotec) were washed with 100 μL of nucleic acid equilibration buffer (Miltenyi Biotec) and then five times with 100 μL of 1x wash buffer. Beads were applied to the column in 100 μL aliquots and washed five times with 100 μL of 1x wash buffer. Columns were demagnetized and beads eluted with two 100 μL washes with 1x wash buffer, and columns were remagnetized. The 200 μL bead solution was combined with RNA sample and rotated at room temperature for 20 min. The sample was then applied to the column in 100 μL aliquots. Columns were washed three times with 400 μL of wash 1 buffer (10 mM Tris-Cl pH 7.4, 6 M urea, 10 mM EDTA) prewarmed to 65°C and then three times with 400 μL wash 2 buffer (10 mM Tris-Cl pH 7.4, 1 M NaCl, 10 mM EDTA). RNA bound to the column was eluted with five washes of 1x wash buffer with 0.1 M dithiothreitol, and then ethanol precipitated with 2 μL of glycoblue.

#### RNA sequencing

Sequencing libraries were prepared using the TruSeq Stranded mRNA Sample preparation kit (Illumina), according to manufacturer’s instruction manual (Rev. E), and sequenced at The Centre for Applied Genomics (SickKids).

#### Computational analysis: read mapping

Libraries were pooled and sequenced on an Illumina HiSeq 2500 to give 50 bp single-end reads. RTA v1.18.54 or later was used for base calling and quality scores, bcl2fastq2 v2.17 or later was used to demultiplex samples and to convert reads to fastq format. Library quality was assessed using FastQC v0.11.5 (http://www.bioinformatics.babraham.ac.uk/projects/fastqc/). Reads were trimmed and clipped for Illumina adaptors using Trimmomatic v0.36 (Bolger et al. 2014) with the following options LEADING:3 TRAILING:3 SLIDINGWINDOW:4:15 MINLEN:36. Reads were aligned to merged reference genomes (hg38 + sacCer3, hg38 + dm6 + sacCer3, hg38 + dm6 + ce10, mm10 + dm6 + sacCer3) obtained using the UCSC Table Browser (Karolchik et al. 2004; Rosenbloom et al. 2015) and kentUtils v302 using STAR version 2.5.2a_modified (Dobin et al. 2012). STAR was invoked with default settings aside from outFilterMultimapNmax 10, outFilterMismatchNoverLmax 0.05, outFilterScoreMinOverLread 0.75, outFilterMatchNminOverLread 0.85, alignIntronMax 1, and outFilterIntronMotifs RemoveNoncanonical.

Mapped reads were quantified using two different methods. First, an in-house R script (v3.2.3) was used to select the longest transcript for every gene, using the GenomicFeatures (Lawrence et al. 2013), rtracklayer (Lawrence et al. 2009), and plyr packages as well as packages from Bioconductor (Gentleman et al. 2004; Huber et al. 2015). Any introns that overlapped with an exon in any other isoform were removed. HTSeq 0.6.1p2 (Anders et al. 2015) was used to count reads mapping to introns and exons. In addition to HTSeq, an in-house intersection method was used, based on tools provided by BEDTools suite v2.26.0 (Quinlan and Hall 2010). Briefly, rather than counting the number of reads mapping to a feature, coverage was determined at the nucleotide level and subsequently averaged at the feature level.

#### Computational analysis: half-life determination

All downstream analyses were performed using in-house R scripts utilizing the follow libraries: scales, plyr, gplots, Hmisc, and limma (Ritchie et al. 2015). Read counts were first filtered to require that each gene had a minimum of one mapped read in all timepoints with five or more mapped reads in at least one time point. Transcriptomic reads were normalized to spike-ins. For transcription inhibition experiments, half-lives were determined by fitting an exponential decay model to normalized data using nonlinear least squares. For 4SU time courses, a bounded growth equation was fit using weighted nonlinear least squares. In DRUID, introns were also used. Introns were filtered such that the mean coverage spanning the intron was 0.5 reads and clustered based on their time-dependent expression profiles using k-means clustering with four clusters. The cluster exhibiting behavior closest to the expected non-increasing time-dependent abundance was used. DRUID is available on GitHub: https://github.com/risslandlab/DRUID

A list of human-mouse orthologs was downloaded from Mouse Genome Informatics (http://www.informatics.jax.org/).

